# Association of physiological variables with football subconcussive head impacts: Why measure?

**DOI:** 10.1101/493700

**Authors:** Megan E. Huibregtse, Steven W. Zonner, Keisuke Ejima, Zachary W. Bevilacqua, Sharlene Newman, Jonathan T. Macy, Keisuke Kawata

## Abstract

Subconcussive head impacts, defined as impacts to the cranium that do not result in clinical symptoms of concussion, are gaining traction as a major public health concern. Researchers begin to suggest subconcussive impact-dependent changes in various neurological measures. However, a contribution of physiological factors such as physical exertion and muscle damage has never been accounted. We conducted a prospective longitudinal study during a high school American football season to examine the association between physiological factors and subconcussive head impact kinematics. Fifteen high-school American football players volunteered in the study. A sensor-installed mouthguard recorded the number of head impacts, peak linear (PLA: *g*) and peak rotational (PRA: rad/s^2^) head accelerations from every practice and game. Serum samples were collected at 12 time points (pre-season baseline, five in-season pre-post games, and post-season) and assessed for the creatine kinase skeletal muscle-specific isoenzyme (CK-MM), as a surrogate for skeletal muscle damage. Physical exertion was estimated in the form of excess post-exercise oxygen consumption (EPOC) from heart rate data captured during five games via a wireless heart rate monitor. A total of 9,700 hits, 214,492 *g*, and 19,885,037 rad/s^2^ were recorded from 15 players across the study period. Mixed-effect regression models indicated that head impact kinematics (frequency, PLA, and PRA) were significantly and positively associated with CK-MM increase, but not with EPOC. There was a significant and positive association between CK-MM and EPOC. These data suggest that skeletal muscle damage effects should be considered when using outcome measures that may have an interaction with muscle damage, including inflammatory biomarkers and vestibular/balance tests.

## Introduction

Repetitive head impacts observed in sports have become one of the most complex public health issues. The majority of head impacts do not induce clinical signs and symptoms of concussion and are often referred as subconcussive head impacts [1]. Despite the individual being asymptomatic, these head impacts have the potential to cause insidious neurological deficits, if sustained repetitively over time [2, 3]. Emerging evidence has postulated that long-term exposure to repetitive subconcussive head impacts is a key predictor for the development of a neurodegenerative pathology called chronic traumatic encephalopathy (CTE) [4, 5]. However, such a cause and effect relationship can only be confirmed by a life-long longitudinal study, which has been deemed infeasible.

A number of studies have been conducted to examine neuronal responses to subconcussive impacts in both acute (several days) and subacute (several weeks) time frames, in addition to the chronic effects from an entire season. Collectively, subconcussive head impacts have shown to induce transient vestibular defect [6], lingering ocular-motor impairment [7, 8], acute and chronic elevations in blood biomarkers of neural injury [9, 10], and pronounced axonal diffusion over the course of a season [3, 11, 12]. A concern has been raised that virtually all reports to date were unable to account for potential confounding effects from vigorous exercise and muscle damage, with some papers listing this issue as a limitation [7, 10, 13, 14] as evidenced by the following quotes, while other papers simply failed to acknowledge it [15-17].

> *“Further validation is needed to correlate systemic biomarkers to repetitive brain impacts, as opposed to the extracranial effects common to an athletic population such as exercise and muscle damage.”*
>
> — – Di Battista et al. 2016 [13]

> *“A lack of biomechanical, neuroimaging, and neuropsychological data limit our ability to determine if the elevations in serum neurofilament-light are a result of axonal damage caused by head impacts or from another source, such as muscle.”*
>
> — – Oliver et al. 2016 [10]

During periods of physical exertion in a hot environment, the core body temperature can exceed 40°C (104°F) [18]. An animal study indicated that when animals sustained traumatic brain injury under hyperthermic conditions, there was greater neuronal cell death in the hippocampus compared to normothermic conditions [19]. In addition to the thermic effect of activity, intense exercise often triggers muscle damage, systemic inflammation, and fatigue, which may influence neurological variables including neurocognitive function [20], blood biomarker [21, 22], and balance [23]. If effects from physical exertion and muscle damage are associated with either subconcussive impact frequency/magnitude or neurological outcome variables, caution is warranted in interpreting the previous reports indicating subconcussive impact-dependent increase in neural injury blood biomarkers [9, 10, 16], neural network disruption [11, 12], and ocular-motor impairment [7]. Conversely, if such an interaction between the physiological factors and subconcussive head impacts does not exist, it is unnecessary to account for physiological factors in the causal inference of head impacts on brain damage.

Given that gold-standard neurological measurements for subclinical neural injury have not yet been established, we instead examined the association between physiological factors and subconcussive head impact kinematics in a prospective longitudinal study of high-school football players during a single season. For head impact kinematics, we employed a sensor-installed Vector mouthguard, whose kinematic accuracy has shown to be superior to helmet, skin patch and headband sensors [24-26], to record frequency and magnitude of head impacts from all practices and games. For physical exertion, we monitored players’ heart rate via wireless chest-strap heart-rate monitor and estimated players’ excess post-exercise oxygen consumption (EPOC). EPOC is a well-accepted measurement in the field of exercise physiology to estimate the degree of “oxygen debt” during exercise, reflecting physiological load and energy metabolism [27]. For muscle damage, we collected blood samples at pre- and post-games from five in-season games and measured serum levels of creatine kinase (CK), particularly a skeletal muscle-specific isoenzyme (CK-MM), which has been considered an indirect surrogate marker of muscle damage [28]. Collectively, we tested our hypotheses that there would be no significant association between head impact kinematics and physiological factors (EPOC and CK-MM), while there would be a significant association between EPOC values and creatine kinase levels.

## Materials and methods

### Participants

Seventeen high school football players at Irvington High School of the National Federation of State High School Associations (NFHS) volunteered for this study. The study was conducted during the 2017 football season, including a preseason physical examination on July 18, 2017, five in-season games (September 22, October 16, 20, and 28, and November 4, 2017), and postseason follow-up (December 1, 2017: Fig 1). None of the 17 players were diagnosed with a concussion during the study period as confirmed by the team athletic trainer and physician. Inclusion criterion included being an active football team member. Exclusion criteria included a history of head and neck injury in the previous 1 year or neurological disorders. Two players actively withdrew within the first month, and their data were not included in the analyses. All participants and their legal guardians gave written informed consent, and the Washington Hospital Healthcare System Institutional Review Board approved the study.

**Fig 1.**
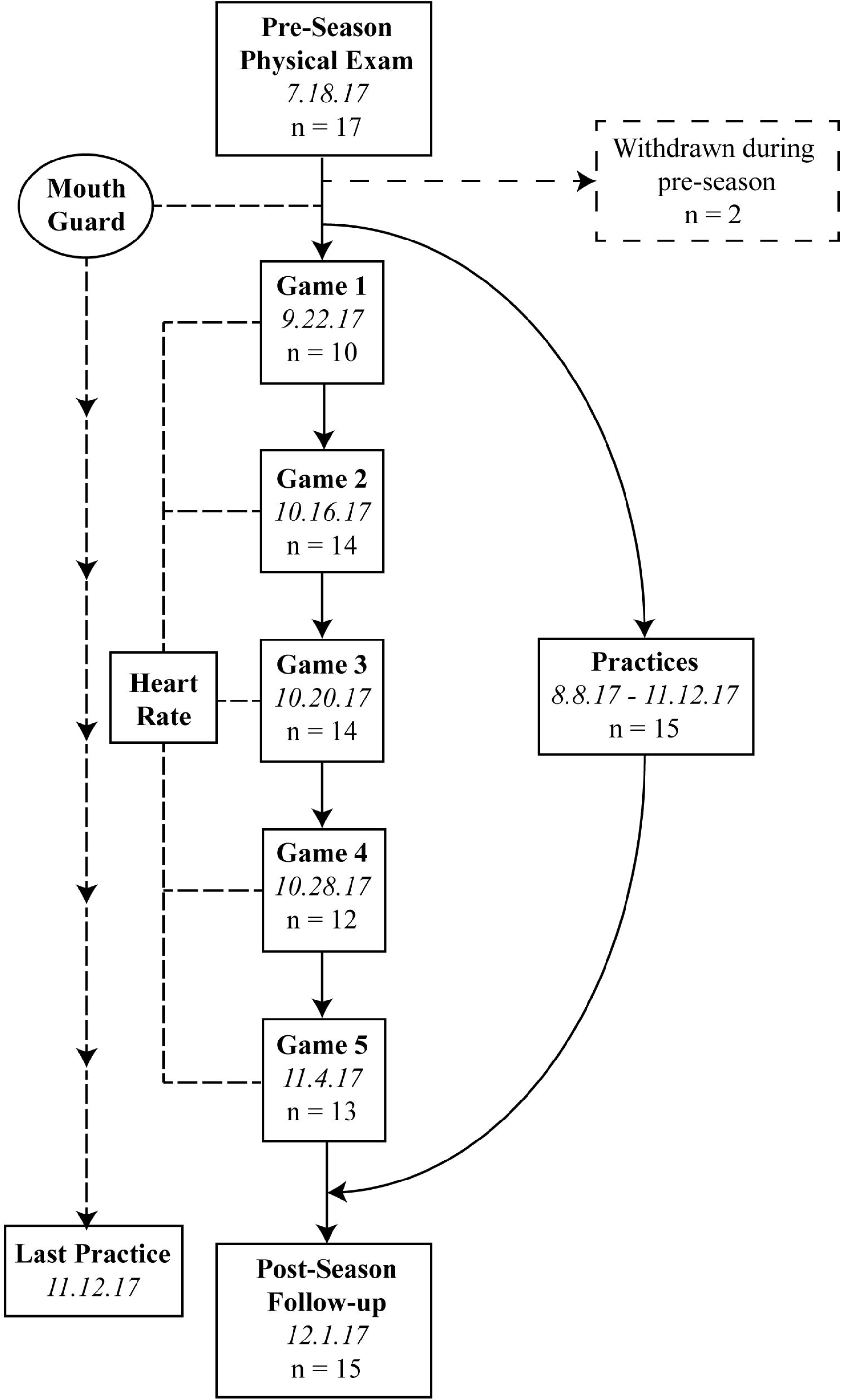
Study Flow Chart.

### Study procedures

During the preseason physical examination, participants were custom-fitted with the Vector mouthguard (Athlete Intelligence, Kirkland, WA) that measured the number of hits and magnitude of head linear and rotational acceleration. Players wore the mouthguard for all practices and games from the beginning of the summer training camp (August 8, 2017) to the end of the season (November 11, 2017). Players were also fitted with a wireless chest-strap heart-rate monitor (Firstbeat Technologies, Jyväskylä, Finland) to record variability of heart rate during the five games. During the preseason baseline assessment, self-reported demographic information (age, height, weight, history of concussion, and years of American football experience) and blood samples were collected. There were no practices or games between the final day of the season (November 11, 2017) and post-season data collection (December 1, 2017).

### Head impact measurement

This study used an instrumented Vector mouthguard for measuring linear and rotational head kinematics during impacts as previously described [7, 9, 26]. The mouthguard employs a triaxial accelerometer (ADXL377, Analog Devices, Norwood, MA) with 200 g maximum per axis to sense linear acceleration. For rotational kinematics, a triaxial gyroscope (L3GD20H, ST Microelectrics, Geneva, Switzerland) was employed. When a preset threshold for a peak linear acceleration (PLA) magnitude exceeded 10.0 g, 16 pre-trigger and 80 post-trigger samples with a standard hit duration of 93.75 milliseconds of all impact data were transmitted wirelessly through the antenna transmitter to the sideline antenna and computer, then stored on a secure internet database. The Vector mouthguard employs an in-mouth sensor to ensure that data acquisition occurs only when the mouthguard is securely fitted in one’s mouth. From raw impact data extracted from the server, the number of hits, PLA, and peak rotational acceleration (PRA) were used for further analyses. Four observations were consistent outliers on the number of hits, sum of PLA, and sum of PRA, exceeding at least 5.5 standard deviations above the mean. These observations were excluded from analysis (<2.5% of all data).

### Blood collection and creatine kinase measurements

Blood samples were obtained from 12 time points. For the five game days, pre-game blood samples were collected four to five hours prior to competition to ensure no effect from pre-game warm up, and post-game blood samples were collected within one hour after the games. At each time point, four milliliters of venous blood samples were collected into red-cap serum vacutainer sterile tubes (BD Bioscience). Blood samples were allowed to clot at room temperature for a minimum of 30 minutes. Serum was separated by centrifugation (1500 x g, 15 min) and stored at -80°C until analysis. CK-MM measurements were performed using sandwich-based enzyme-linked immunosorbent assay (ELISA) kits (LifeSpan Biosciences Inc., Seattle, WA). The lowest detection limit of the assay is 0.94 ng/mL, and the assay covers a concentration range up to 100 ng/mL with a typical intra-assay precision of <6.8% and an inter-assay precision of <5.3%. One hundred microliters of serum samples were loaded in duplicate into the ELISA plate. Fluorescent signals measured by a micro-plate reader (BioTek EL800, Winooski, VT) were converted into ng/mL as per standard curve concentrations. The experimenter performing the assay was blinded from subject information.

### Physiological load measure

EPOC can be estimated by VO_2_ levels and duration of the activity: EPOC (ml/kg) = (mean VO_2 recovery_ x time _recovery_) – (VO_2baseline_ x time _recovery_). To maximize clinical application, instead of VO_2_, researchers have validated an EPOC measurement using %HR_maximum_, time, and heart rate variability, which yielded a substantial agreement with VO_2_-derived EPOC (range of R^2^: 0.79 – 0.87) [29, 30]. Players in this study wore chest-strap HR monitors from pre-game warm up until approximately 1h post-game. Peak EPOC values were not utilized because EPOC declines over time and not necessarily in a linear fashion. Thus, the mean EPOC value from each game was assessed for each player using Firstbeat PRO heartbeat analysis software version 1.4.1 and included in analyses.

### Statistical analysis

A repeated measures analysis of variance was used to test whether EPOC and CK-MM varied across five games, as well as between pre- and post-games for CK-MM levels. Mixed-effect regression modeling (MRM) was performed using physiological factors (EPOC and CK-MM changes between pre- and post-games) and three head impact kinematics (PLA, PRA and head impact counts) in each game as primary outcomes. All the models were adjusted by cumulative frequency and magnitude of head impacts up to each game (from previous games and practices). Pre-season baseline CK-MM level and a pre-game CK-MM level were additionally adjusted for in the models that used CK-MM change as an outcome. Furthermore, to examine whether physical exertion and muscle damage parameters were correlated with one another, the potential correlation between EPOC and acute changes in CK-MM levels between pre- and post-games was assessed using MRM where CK-MM change as the outcome and EPOC as a primary predictor. The model was adjusted by pre-season baseline CK-MM level and pre-game CK-MM level. All analyses were conducted using statistical software R (version 3.4.1) with package ‘nlme’.

## Results

### Demographics and head impact kinematics

A total of 9,700 hits, 214,492 *g*, and 19,885,037 rad/s^2^ were recorded from 15 players during pre-season training camp, in-season, and post-season stages, collectively. Demographic characteristics and median values of impact kinematics are summarized in Table 1. Please refer to Supplementary Table 1 for the impact data from each game.

**Table 1.** **Demographics and head impact kinematics.**

| **Table 1. Demographics and head impact kinematics.** |  |
| --- | --- |
| **Variables** | **Team (n=15)** |
| **Demographics, M (SD)** |  |
| Age, y | 16.4 (0.5) |
| Body Mass Index, kg/m<sup>2</sup> | 28.0 (4.0) |
| Years Football Experience | 3.13 (1.5) |
| Number of previous concussions | 0.47 (0.6) |
| 0 | 9* |
| 1 | 5* |
| 2 | 1* |
| **Position, n (%)** |  |
| Linemen (OL, DL) | 6 (37.5) |
| Linebacker, Tight End, | 2 (18.7) |
| Skill Players (WR, DB, RB) | 7 (43.8) |
| **Impact Kinematics for Season, median (IQR)<sup>†</sup>** |  |
| Total number of hits | 596 (361.5 – 981.5) |
| Sum of PLA, g | 11,907 (7,644 – 21,014) |
| Sum of PRA, rad/sec<sup>2</sup> | 1,202,758 (637,515 – 210,0871) |
| Note: IQR, interquartile range. SD, standard deviation. OL, offensive lineman. DL, defensive lineman. WR, wide receiver. DB, defensive back. RB, running back. PLA, peak linear acceleration. PRA, peak rotational acceleration.<br>*number of players, <sup>†</sup>see supplemental table 1 for impact kinematics from individual games |  |

### Patterns of EPOC and creatine kinase changes across five games

In the overall sample, repeated measures ANOVA indicated that there was no difference in EPOC between games, as illustrated by a statistically non-significant time effect, F(1,46)=1.213, p=0.277. Likewise, there was no significant change in serum CK-MM levels across study time points, F(1,48)=0.495, p=0.485 (Table 2).

**Table 2.** Means and standard deviations of EPOC and CK-MM across five football games.

|  | Game 1 |  | Game 2 |  | Game 3 |  | Game 4 |  | Game 5 |  |
| --- | --- | --- | --- | --- | --- | --- | --- | --- | --- | --- |
| EPOC<sup>a</sup> | 17.25 (18.85) |  | 13.45 (12.81) |  | 28.43 (22.74) |  | 25.98 (33.73) |  | 14.57 (9.724) |  |
| CK-MM<sup>b</sup> | Pre | Post | Pre | Post | Pre | Post | Pre | Post | Pre | Post |
|  | 7.893<br>(6.641) | 9.184<br>(5.815) | 12.47<br>(6.000) | 13.39<br>(7.362) | 8.010<br>(5.745) | 9.879<br>(5.133) | 7.338<br>(3.217) | 9.822<br>(4.360) | 9.195<br>(4.330) | 12.06<br>(7.106) |

Note: EPOC, excess post-exercise oxygen consumption; CK-MM, creatine kinase muscle specific isoenzyme.
<sup>a</sup>mL/kg, expressed as mean (SD)
<sup>b</sup>ng/mL, expressed as mean (SD)

### Association between physiological factors and head impact kinematics

Mixed-effect regression models indicated that head impact kinematics (frequency, PLA, PRA) were significantly positively associated with CK-MM increase (Fig 2), but not with EPOC (Table 3). For example, the model estimated that CK-MM at post-game would increase by 0.06 ng/mL (SE=0.03) for each additional head impact sustained during a game. There was a significant positive association between EPOC and CK-MM increase (β=0.07, SE=0.025, P=0.009; Fig 3).

**Table 3.** Results from mixed-effects regression models.

|  | Increase in CK-MM (ng/mL) |  |
| --- | --- | --- |
|  | Estimate (SE) | P-value |
| **In-game head impact count (times)** | 0.06 (0.029) | 0.041 |
| **In-game PLA (g)** | 0.003 (0.001) | 0.031 |
| **In-game PRA (rad/s<sup>2</sup>)/1000** | 0.029 (0.013) | 0.029 |
|  | EPOC (mL/kg) |  |
|  | Estimate (SE) | P-value |
| **In-game head impact count (times)** | 0.073 (0.17) | 0.672 |
| **In-game PLA (g)** | 0.001 (0.007) | 0.882 |
| **In-game PRA (rad/s<sup>2</sup>)/1000** | 0.008 (0.076) | 0.918 |
| Note. PLA, peak linear acceleration; PRA, peak rotational acceleration; CK-MM, creatine kinase skeletal muscle isoenzyme; EPOC, excess post-exercise oxygen consumption. |  |  |

**Fig 2.**
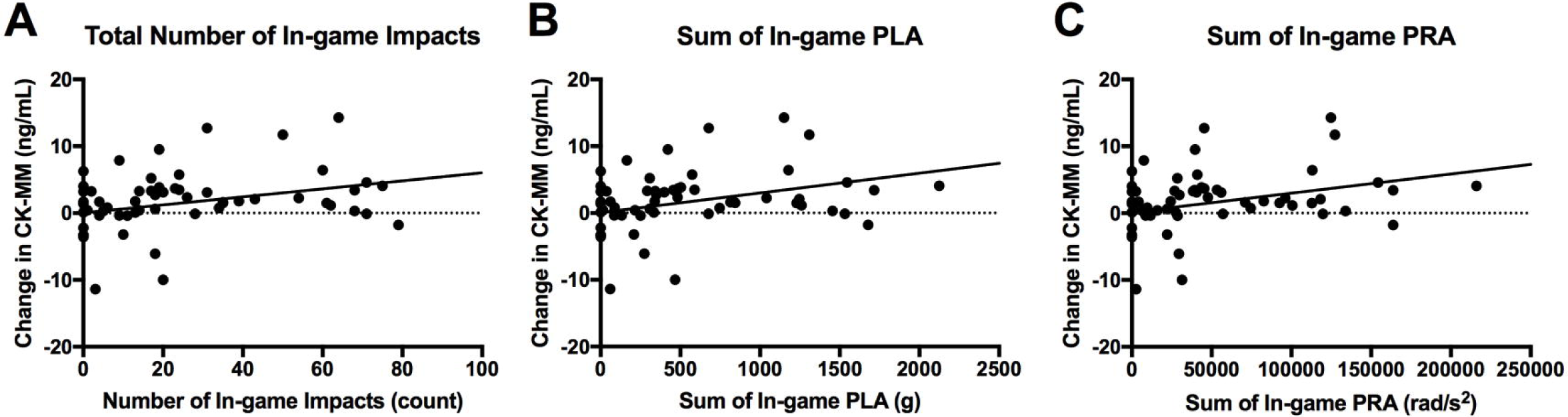
Association between Change in CK-MM and Head Impact Kinematics. The change in CK-MM from pre- to post-game was significantly associated with the frequency of hits (Panel A; β=0.06, SE=0.029, P=0.041), the sum of peak linear acceleration sustained during the game (Panel B; β=0.003, SE=0.001, P=0.031), and the sum of the peak rotational acceleration of impacts sustained during the game (Panel C; β=0.029, SE=0.013, P=0.029).

**Fig 3.**
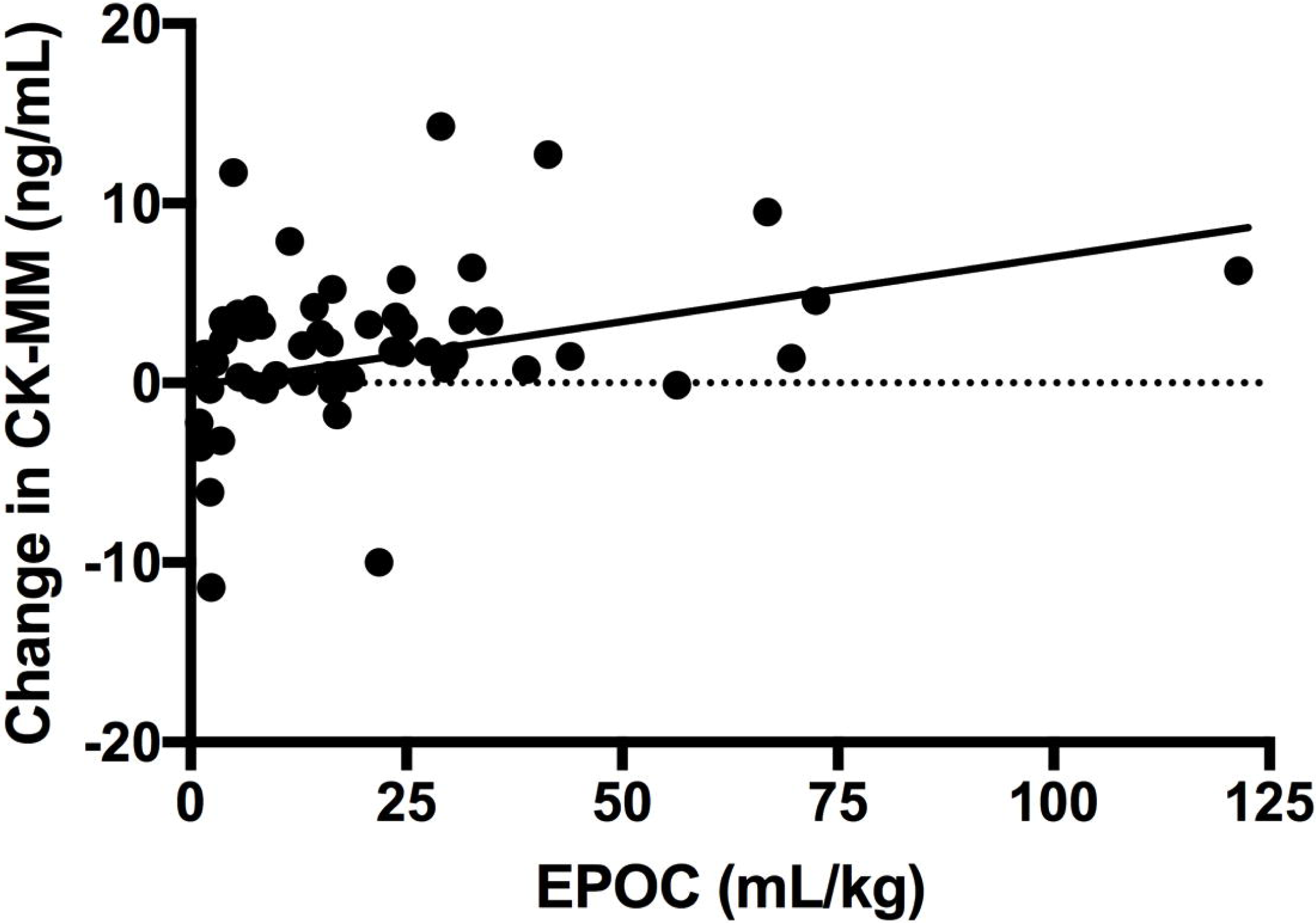
Association between Change in CK-MM and EPOC. There was a significant positive association between the EPOC and the change in CK-MM from pre- to post-game (β=0.07, SE=0.025, P=0.009).

## Discussion

In the present study, we assessed two of the most practical physiological variables, physical exertion and muscle damage, and examined their association with head impact frequency and magnitude sustained during football games. To the best of our knowledge, this is the first and most comprehensive clinical study to suggest the potential tri-linkage among head impact kinematics, muscle damage, and physical exertion. There were three chief findings from the present study: 1) physical exertion levels as measured by EPOC were not associated with frequency and magnitude of head impacts sustained during games, 2) muscle damage as assessed by CK-MM was significantly and positively associated with head impact frequency and magnitude, and 3) there was a significant positive association between muscle damage and physical exertion measures.

The most meaningful implication of this study is that it may be unnecessary to account for physical exertion effects in subconcussion studies. This information is significant for existing literature that failed to measure physical exertion variables [7, 10, 13, 14, 16, 17, 31], as well as planning for future studies, given the limited feasibility of heart rate monitor usage by athletes. Contrarily, we highly encourage researchers to account for muscle damage effects, especially for studies using outcome variables that can be influenced by muscle damage, such as inflammatory markers, brain-enriched factors that are expressed, albeit low level, within skeletal muscle (i.e., S100B), and balance assessment. This recommendation is supported by our novel and unexpected finding that there was a significant positive association between serum CK-MM levels and head impact kinematics, which necessitates a brief review of the mechanism by which CK-MM is released into the bloodstream.

CK-MM is constitutively localized to the M-line of sarcomeres in muscle fibers and facilitates the regeneration of adenosine triphosphate (ATP) for muscle fibers [32]. Traumatic stress and/or mechanical stress to the sarcomere can disrupt the contractile apparatus, muscle cytoskeleton, and myofibrillary enzymes such as CK-MM. The dislodged CK-MM can translocate into the extracellular space and then to the bloodstream [33]. Resistance exercise, particularly eccentric action like biceps curls and leg presses, has consistently shown to increase serum levels of CK-MM. Ehlers et al. reported that a two-a-day practice in college American football players elevated serum CK levels 15-fold [34], supporting the strenuous nature of American football and its highest rate of musculoskeletal injury of any team sport [35]. Although our sample showed lesser magnitudes of acute CK-MM increases than the Ehlers et al. [34], our data indicate that those who sustained greater frequency and magnitude of head impacts are likely to experience greater muscle damage as reflected by increased serum CK-MM levels. Some may claim that this observation is as expected, given that the more plays in which an athlete participates, the more likely he/she triggers explosive muscle contraction to block, sprint, and/or tackle, resulting in a greater chance of sustaining subconcussive head impacts. Playing positions may provide a clearer interpretation of the data, since play style and frequency of head impacts [1] are largely dependent on players’ position. However, due to small sample size and some subjects participating in both defensive and offensive plays, we were unable to conduct such a position dependent analysis.

A perplexing aspect of the results was that the head impact kinematics were significantly associated with CK-MM levels but not with EPOC values, despite these two physiological variables being significantly correlated with one another. Similar to CK-MM, EPOC is particularly sensitive to anaerobic exercise, such that a high-intensity resistance exercise has shown to exhibit the greatest increase in EPOC, whereas smaller increases in EPOC have been documented after a long duration of aerobic exercise [27]. American football is comprised of frequent bouts of anaerobic exercise, supporting the appropriate use of EPOC in understanding physical exertion load. A new hypothesis generated by this independent relationship between EPOC and head impact kinematics is that an EPOC value is relatively consistent across plays, while head impact occurrence is subject to several factors. Previous research has shown that impact incidence among American football players is skewed, with lineman (defensive and offensive) and linebackers recording the greatest frequency of head impacts during both games and practices [36]. These findings underpin common speculation that “the box” encompassing the aforementioned positions subjects players to frequent head insults. Additionally, the primary emphasis of American football is to tackle the individual who possesses the football, therefore the risk of head impact seemingly shifts drastically towards the ball carrier, relieving others of harm. Lastly, regardless of player position (inside or outside the box) or frequency of possessing the football, it is understood that all players engage in high intensity anaerobic activity each play, thus creating a uniform rate of physiological demand that is independent of head impact kinematics. It is important to note that our available data cannot rigorously test this speculation, which warrants further investigation to include other physiological factors such as running velocity, blood flow velocity, and PaO_2_.

The impact of these results is limited by several factors. The study cohort was comprised of a small number of male, adolescent football players. Therefore, generalizability of the results should only be limited to the football cohort at best. It remains unclear whether a similar observation may be present in females and in other contact sports (i.e., soccer, ice hockey, boxing). Importantly, this is the first field study to measure and comprehensibly test the relationship between physiological variables and subconcussive head impacts across an entire high school season of American football.

## Conclusions

We sought to determine the relationship between two physiological variables, CK-MM and EPOC, and subconcussive head impacts, as the effect of exercise is often not accounted for or is cited as an unknown quantity in studies of subconcussive head impacts. Data suggest that the variable of physical exertion (EPOC) was independent from subconcussive head impacts. However, we identified that serum levels of CK-MM, reflecting muscle damage, raised concomitantly with the frequency and magnitude of head impacts sustained during football games. This study proved the feasibility of such physiological measurements. However, the data indicate that it may be unnecessary to account for physical exertion effect in subconcussion studies. Conversely, future study designs should include measures of muscular damage in order to distinguish the effects between muscle damage and subconcussive head impacts.

## Supporting information

## Acknowledgments

Authors would like to thank Ms. Ciara Fulgar and Ms. Carmen Charleston for their assistance in data collection. Authors also would like to thank Ms. Michelle Stone, Ms. Lamiya Sheikh, Ms. Kimberly Hartz for their administrative assistance. Additionally, authors would like to acknowledge Ms. Niyati Patel, CMA CPC, Mr. Jacob Geier, Ms. Yesenia Galea, MA, CP-II, Ms. Renae Mullins, CMA, Ms. Vanquynh Pham, RN, FNP, Ms. Annalisa Rocha, Mr. Baldeep Bajwa, CPC, Ms. Leanne Saud, Ms. Cassandra Manuel, RN, and Ms. Laura Constantine in administration and blood sample collection.

## Supporting information

**S1 Table. Impact kinematics for individual games.**

