## Supplementary material for "Association of physiological variables with football subconcussive head impacts: Why measure?"

| *Supplemental Table 1. Head impact kinematics of each game* | | | | | |
| --- | --- | --- | --- | --- | --- |
|  | Game 1 | Game 2 | Game 3 | Game 4 | Game 5 |
| Hits | 14.0  (5.0–21.5) | 18.5  (8.5–29.3) | 20.0  (17.5–48.0) | 31.0  (14.0–37.0) | 50.0  (23.5–66.0) |
| PLA (*g*) | 304.0  (85.5–369.3) | 380.5  (203.0–652.0) | 421.0  (300.0–1,037.0) | 482.0  (268.5–822.0) | 1,150.0  (566.5–1,382.0) |
| PRA (rad/s^2^) | 20,181.7  (7,861.5–36,250.1) | 35,320.7  (19,961.2–45,316.0) | 39,711.4  (24,510.4–96,604.2) | 56,045.1  (23,243.0–78,631.0) | 118,255.1  (48,252.6–130,624.0) |
| Note: Impact kinematics data are expressed as median (IQR). IQR, interquartile range. PLA, peak linear acceleration. PRA, peak rotational acceleration. | | | | | |
